# Defining the Growth Stages of Mungbean (*Vigna radiata* L.) using the BBCH Scale

**DOI:** 10.1101/2025.11.09.687504

**Authors:** Venkata Naresh Boddepalli, Arti Singh

## Abstract

Mungbean (*Vigna radiata* L.) is a protein-rich food grain legume that fixes nitrogen and serves as a potential crop for the plant-based protein industry. It can fit in multiple cropping systems as a short-duration and low-water requirement crop. Phenological growth stages of mungbean are described here for the first time according to the BBCH (Biologische Bundesanstalt, Bundessortenamt, and Chemische Industrie) scale. We developed an extended BBCH chart with a two-digit scale to identify different phenological growth stages of mungbean. This study documented and illustrated the key growth stages of mungbean from germination to maturity, correlating them with the number of days after planting and accumulated growing degree days (GDD). It outlines the principal growth stages (PS) as follows: germination (PS-0), leaf development (PS-1), stem elongation (PS-3), flower bud development (PS-5), flowering (PS-6), fruit development (PS-7), ripening (PS-8), and senescence (PS-9). This chart can help track the key mungbean growth and developmental stages. Most importantly, this BBCH growth stages chart fills a gap in the literature by providing a practical tool for researchers and growers to standardize crop management practices, develop simulation models, and assess performance under various environmental conditions. The standardized scale enhances the precision of timing agricultural interventions, ultimately supporting improved mungbean production in the region.

## INTRODUCTION

The global population is estimated to reach 9.7 billion by 2050 (United Nations Publications, 2023), intensifying concerns about food security and the need for sustainable increases in agricultural productivity (Oluwole et al., 2023). To address these challenges, diversified cropping systems have emerged as a promising strategy, offering both environmental and nutritional benefits (Douyon et al., 2022; Yang et al., 2024). Among these systems, the integration of legumes into crop rotations has gained significant attention due to their ability to increase yields by 38%, reduce greenhouse gas emissions, and improve soil health through biological nitrogen fixation (Kebede, 2021; Mesfin et al., 2023; Yang et al., 2024). In addition, legume crops are rich in proteins, carbohydrates, essential amino acids, micronutrients, and vitamins. They are able to play a crucial role in combating malnutrition, particularly in developing countries where access to animal-based proteins is limited (Langyan et al., 2022; Samal et al., 2023). The interest in plant-based protein sources also aligns with sustainable dietary practices, also their contribution to reducing the environmental footprint of food production (Langyan et al., 2022; Viroli et al., 2023). As the demand for sustainable and nutritious food sources grows, a practical strategy is to integrate diverse legumes into global agricultural systems to enhance food security, improve nutrition, and promote environmental sustainability (Lisciani et al., 2024).

Mungbean (*Vigna radiata* L.), a food grain legume that originated in the Indo-Burma region, is widely cultivated in Asian countries and is gaining popularity in other regions, including Africa, Australia, and parts of the United States (Nair & Schreinemachers, 2020; Somta et al., 2022). Its increasing adoption is driven by its high nutritional value, particularly its digestible proteins (24–28% on a dry weight basis), minerals, vitamins, and essential micronutrients, as its ability to fix atmospheric nitrogen, and its economic importance (Liang et al., 2020; USDA Natural Resources Conservation Service, 2024). Mungbean adaptability to warm climates and its short duration (65-75 days after planting), coupled with low input requirement, makes it a versatile crop that can be integrated into various agricultural systems, such as monoculture, double cropping, or as a cover crop (Jaya et al., 2021; Koyejo et al., 2021; Batzer et al., 2022b). In the Midwestern United States, the existing soybean infrastructure can be readily adapted for mungbean cultivation with minimal adjustments, positioning it as a viable alternative crop for farmers in the region(Batzer et al., 2022a).

Understanding the phenological growth stages of mungbean is essential for effective production management. Such knowledge enables the synchronization of management practices with critical growth stages, crop monitoring, optimizing crop performance, and resilience (You et al., 2022; Liu et al., 2023). A standardized framework for describing plant growth stages is indispensable for accurate crop management, modeling, and evaluation under various environmental conditions and input responses (Dambreville et al., 2015; Wang et al., 2024). The BBCH scale is widely accepted for this purpose, providing a universal system to describe and compare growth stages across crop species (Awachare & Upreti, 2020; Taghavi et al., 2022).

Though mungbean is globally cultivated on over 7 million hectares (Nair & Schreinemachers, 2020; Somta et al., 2022), a standardized growth stage chart, according to the BBCH scale, remains unavailable. This can be a caveat for researchers, agronomists, and farmers to optimize management practices, develop accurate crop models, and assess performance under diverse conditions. To address this gap, we proposed this study with a twofold aim: 1) to define phenological growth stages according to the two-digit BBCH scale and 2) to record the number of days after planting and calculate the growing degree days required for each growth stage in the Midwest region of the United States.

## 2. MATERIALS AND METHODS

### 2.1. Experimental location

This study was carried out over two growing seasons (2023 and 2024) at the Iowa State University Research Farm on Agricultural Engineering and Agronomy (AEA) near Boone, Iowa, USA. In 2023, planting occurred on 1 June at the AEA farm (42°01’14.9” N, 93°46’05.1” W). In 2024, planting took place on 2 June at two locations: the AEA farm (42°01’08.0” N, 93°46’18.4” W) and the Bruner farm (42°00’51.5” N, 93°43’52.2” W). All sites were characterized by silty clay loam soils typical of the Des Moines Lobe landform. The previous crop at all locations was corn.

### 2.2. Plant material and experimental design

Two mungbean cultivars, ISU Mung G1 and ISU Mung G2 (Table 1), disclosed by the Iowa State University (ISU) were utilized for this study. These two lines were developed by the ISU breeding program through a modified-bulk method involving bi-parental crosses for the Midwestern growing conditions. The seed of each cultivar was planted in a plot with 10-foot rows, spaced 30 inches apart with 7.5-inch intervals at a depth of 1 inch, using a Randomized Complete Block Design (RCBD) with five replications at each location. Standard agronomic practices were followed during the experiment in these locations.

**Table 1:** Key features of the two cultivars utilized in this study.

| **Trait** | **ISU mung G1** | **ISU mung G2** |
| --- | --- | --- |
| Origin | ISU, Iowa | ISU, Iowa |
| Pedigree | AVMU0201 x PI 425513 | AVMU0201 x PI 377167 |
| Growth Habit | Erect | Erect |
| Days to First Flower | 39 | 41 |
| Days to 80% Mature Pods | 68 | 67 |
| 1000 Seed Weight (g) | 39.3 | 34.1 |
| Seed % Protein (dry basis) | 24.19 | 24.25 |
| Plant Height (at maturity) (cm) | 49.8 | 49.8 |
| Seed Texture | Matte | Shiny |
| Seed Color | Green | Green |

### 2.3. Weather data

Daily weather data, including maximum temperature (Tmax), minimum temperature (Tmin), relative humidity, and precipitation measurements, were obtained from the Iowa Environmental Mesonet website (Herzmann, n.d.), collected from an on-site weather station (Table S1). Figure 1 illustrates the average weekly weather parameters, mainly minimum and maximum air temperatures and precipitation observed during the study for each month and year.

**Figure 1:**
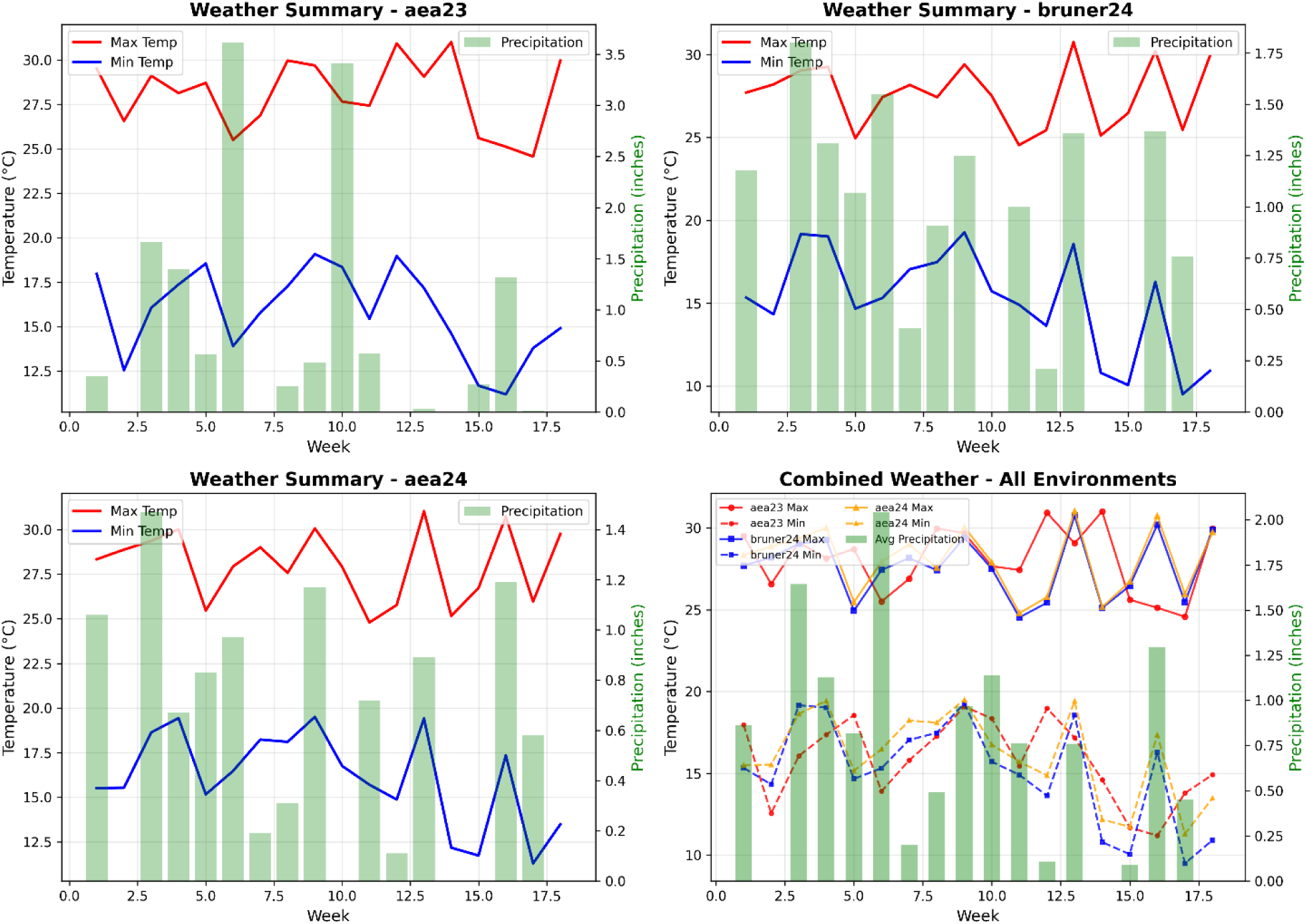
Weekly weather patterns across experimental environments during 2023-24 growing seasons. Individual panels location-year specific weather conditions for A). Agricultural Engineering and Agronomy (AEA) Research Farm 2023, B) Bruner Farm 2024, C) Agricultural Engineering and Agronomy (AEA) Research Farm 2024, and D) depict the comparative weather patterns across three environments.

### 2.4. Growth Stages data

Development of mungbean phenological stages was assessed using the two-digit BBCH coding system following international standards for beans (Hack et al., 1992; Meier, 2018). The BBCH principal stages assigned were: principal stage 0 (germination: VG-VE, BBCH 00-08), principal stage 1 (leaf development: VC-V2, BBCH 09-12), principal stage 3 (stem elongation: V3-V5, BBCH 30-33), principal stage 5 (flower bud development: V6, BBCH 51), principal stage 6 (flowering: R1, BBCH 60), principal stage 7 (fruit development: R2-R4, BBCH 71-79), principal stage 8 (ripening: R5-R7, BBCH 81-89), and principal stage 9 (senescence: BBCH 97-99). Observations of the phenological stages (Table S2) were recorded during the early morning hours, every two days or weekly, depending on the stage of development. Images of the growth stages were captured using a Nikon D3500 DSLR camera.

### 2.5. Growing degree days (GDD) calculation

Growing degree days (GDD, °C) were calculated by subtracting the base temperature (T_b_ = 10^0^C) from the average of the maximum and minimum temperatures recorded each day (Equation 1) (McMaster & Wilhelm, 1997). The cumulative GDD was calculated from the date of planting for each phenological stage.

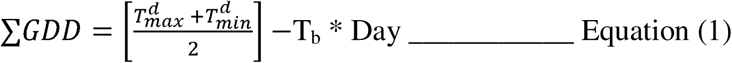

The maximum air temperature (*T^d^_max_*, °C) and minimum air temperature (*T^d^_min_*, °C) are used along with the base temperature (T_b_, °C) to calculate the accumulated thermal time (TTA). The cumulative thermal time (TTA) throughout the crop cycle, as well as for each phenological stage, was calculated by combining the daily thermal time (TTd) required to reach the phenological stage under evaluation, as shown in Equation (2):

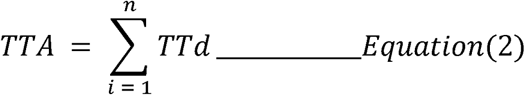

Where n is the number of days required to complete the entire or part of the phenological cycle of the crop. A similar method of calculations was followed by Cavalcante et al. (2020).

### 2.6. Data analysis

Descriptive statistics for DAP and GDD were calculated to determine the mean and standard error for each BBCH stage across environments and cultivars (Pinzón-Sandoval et al., 2024). A linear mixed model: Response ∼ Cultivar + (1|Site) + (1|Year) + (1|Site:Year:Rep) was utilized for analyzing DAP and GDD, where cultivar was treated as a fixed effect, and site, year, and replicate nested within site-year were treated as random effects. Model residuals were checked using the Shapiro-Wilk test and Q-Q plots for normality. Pairwise differences between site-years were calculated using Tukey’s HSD adjustment for multiple comparisons. Correlations between monthly maximum temperature (T_max_), maximum temperature (T_min_), precipitation, and both GDD (°C) and DAP were estimated using Pearson’s correlation coefficient (r) across site-years. Statistical significance was set at α = 0.05. Statistical analyses were performed in Python 3.x using pandas, numpy, and matplotlib packages.

### 2.7. Growth stages sketch

Mungbean growth stages were illustrated by a professional designer to accurately depict each BBCH developmental phase. These illustrations were developed on the basis of comprehensive field observations and photographic documentation of live plant specimens at each growth stage. The digital illustrations were generated using Adobe Photoshop for sketching and coloring, and Adobe Illustrator for final layout and output, with drawing executed using a Wacom graphics tablet. After multiple review sessions throughout the sketching process, the final sketches were reviewed for botanical accuracy and consistency with international standards and then assembled into a comprehensive visual reference guide for mungbean growth stages with V/R code and corresponding days after planting (Figure 6).

## 3 RESULTS

The phenological stages of the mungbean (Table 1) were observed in relation to the chronological time (days after planting), and the thermal time (cumulative degree days) for each stage were depicted in Figures 1-3, and the V/R stage alignment description was presented in Table 2 with DAP and GGD and in Table S3.

**Table 2:**
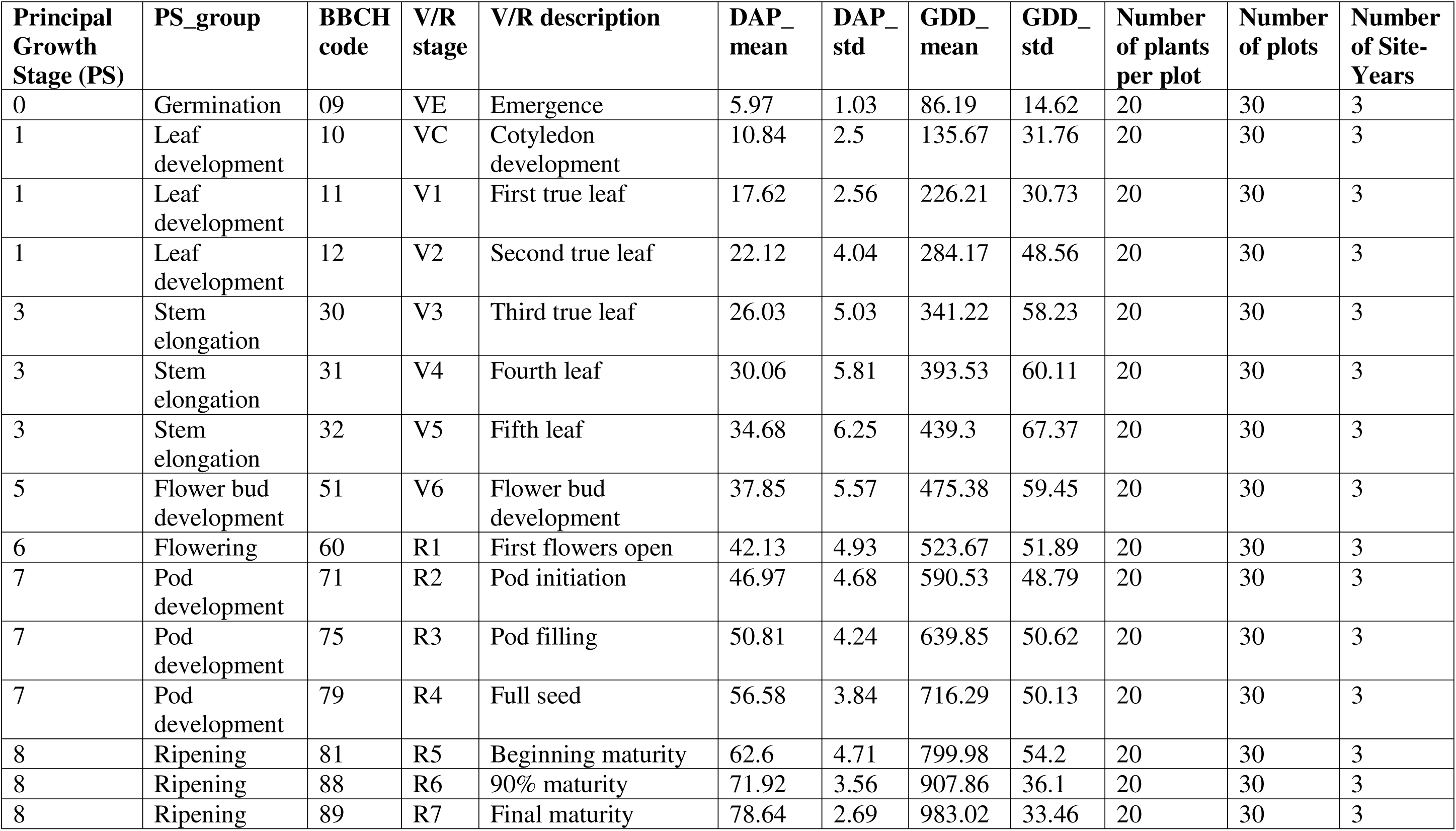
Principal growth stage (PS) and their corresponding vegetative/reproductive (V/R) stages with descriptions for mungbean phenological stages and their mean stage (DAP) and thermal (GDD) timings along with their standard deviations(std).

**Figure 2.**
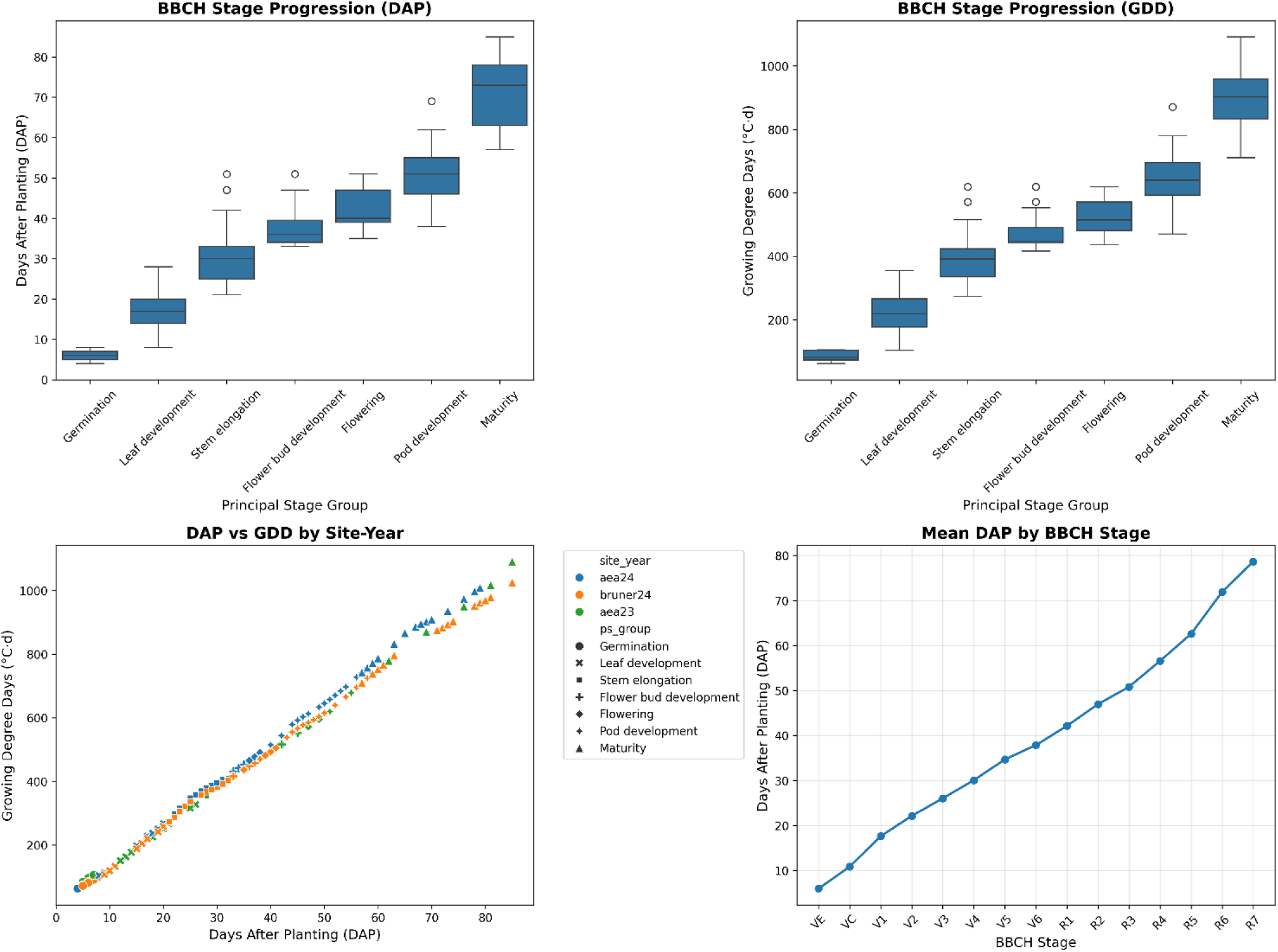
Mungbean principal growth stages progression across multiple site-years. A). DAP distribution box plots with specific BBCH stages listed. B). Accumulation of GDD with the specified base temperature C) Relationship between DAP and GDD showing environmental interactions.

### 3.1. Principal growth stage 0: Germination stage

The germination stage, including dry seed (BBCH 00) that starts imbibing water (BBCH 01). After the complete imbibition stage (BBCH 03), the radicle emergence starts from the seed (BBCH 05), followed by the emergence of the hypocotyl with cotyledons split from the seed coat (BBCH 07), and finally, the hypocotyl reaches the soil surface with a visible arch (BBCH 08), a described in the BBCH scale, as shown in Figure 3.

**Figure 3:**
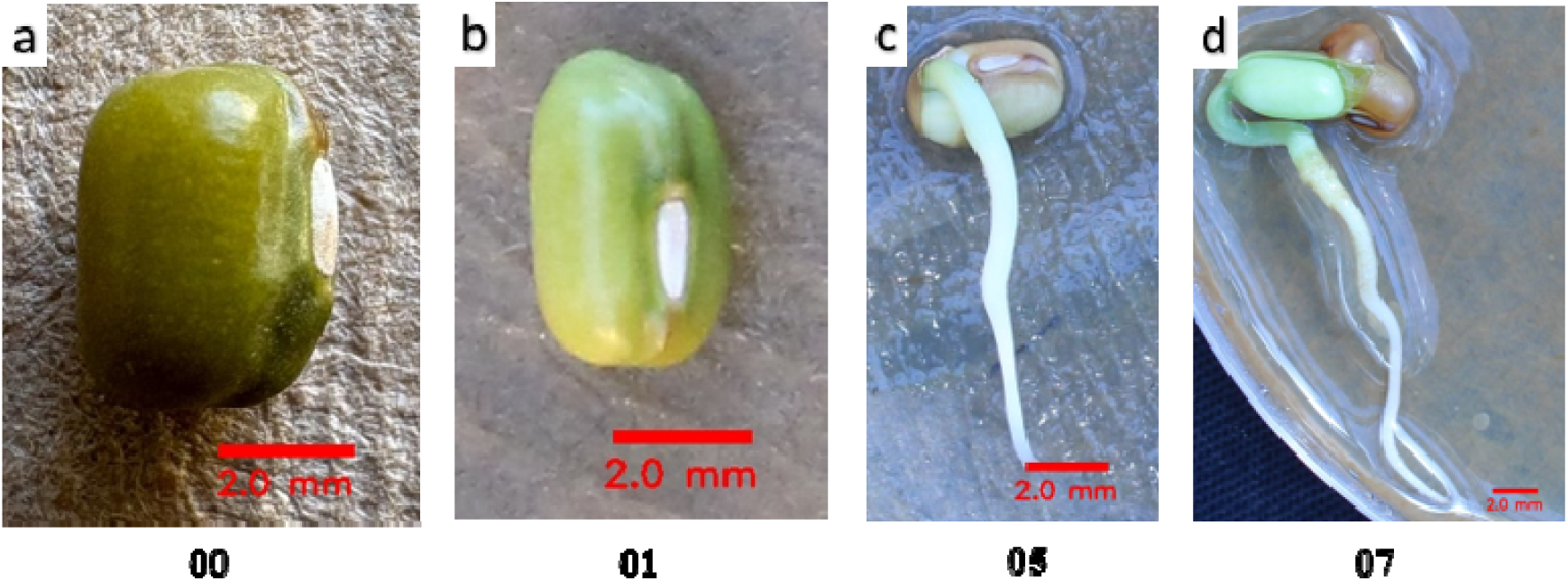
Mungbean germination stage progression according to the BBCH scale. a) dry seed b) starting of the imbibition c) radicle emergence from the seed d) emergence of hypocotyl with split cotyledon, which is ready to crack the soil surface.

#### BBCH 09: VE stage

The emergence stage (VE) occurs when the unifoliate leaves unfurl from the cotyledons above the soil surface, reaching an accumulated thermal unit of 86 °C.d, at 6 day after planting.

### 3.2. Principal growth stage 1: Leaf development

#### BBCH 10: VC stage

This was the beginning of the leaf developmental stage, where the unifoliate leaves fully expand. The edges of the leaves did not touch and were positioned opposite each other on the stem. The development of these leaves required around 188 °C.d and was completed in 10 days after planting.

#### BBCH 11: V1 Stage

The First trifoliate leaves attached to the first node immediately above the unifoliate leaves are fully expanded and flat. Meanwhile, the second trifoliate leaf at the upper node begins to roll. This stage is marked by an accumulated thermal unit of 284 °C.d, reached approximately 17 days after planting. The progression continues until the next significant growth stage.

#### BBCH 12: V2 Stage

The second trifoliate leaf at the second node becomes fully expanded and flat, while the third trifoliate leaf at the upper node starts unrolling. This stage requires an accumulated thermal unit of 284 °C.d, which occurs about 22 days after planting.

### 3.3. Principal Growth Stage 3: Stem elongation

Stem elongation starts at the third leaf stage of the plant, while trifoliate leaves continue their expansion on different nodes. This indicates the transition from leaf development to stem elongation.

#### BBCH 30: V3 Stage

During this stage, the third trifoliate leaf attached to the third node is fully expanded and flat, while the fourth trifoliate leaf at the upper node begins to unroll. The thermal accumulation for this stage is 341 °C.d, reaching approximately 26 days after planting.

#### BBCH 31: V4 Stage

The fourth trifoliate leaf attached to the fourth node is fully expanded and flat, while the fourth trifoliate leaf at the upper node begins to unroll. This stage is marked by an accumulated thermal unit of 393 °C.d, occurring around 30 days after planting.

#### BBCH 32: V5 Stage

During the V5 stage, the fifth trifoliate leaf attached to the fifth node is fully expanded and flat, while the sixth trifoliate leaf at the upper node starts to unroll. This stage is characterized by an accumulated thermal unit of 439 °C.d, approximately 34 days after planting.

### 3.4. Principal Growth Stage 5: Flower bud development

#### BBCH 51: V6 Stage

At stage V6, the sixth trifoliate leaf attached to the sixth node is fully expanded and flat, and the seventh trifoliate leaf at the upper node begins unrolling. The inflorescence developed on the axis of the internodes will start bearing the flower buds. This stage is marked by an accumulated thermal unit of 475 °C.d, occurring approximately 38 days after planting. At this stage, the plants started producing flower buds.

### 3.5. Principal Growth Stage 6: Flowering

The flowers with pale yellow petals start opening, indicating the plant’s transition to the reproductive stage from the vegetative stage (Figure 4 and 5).

**Figure 4.**
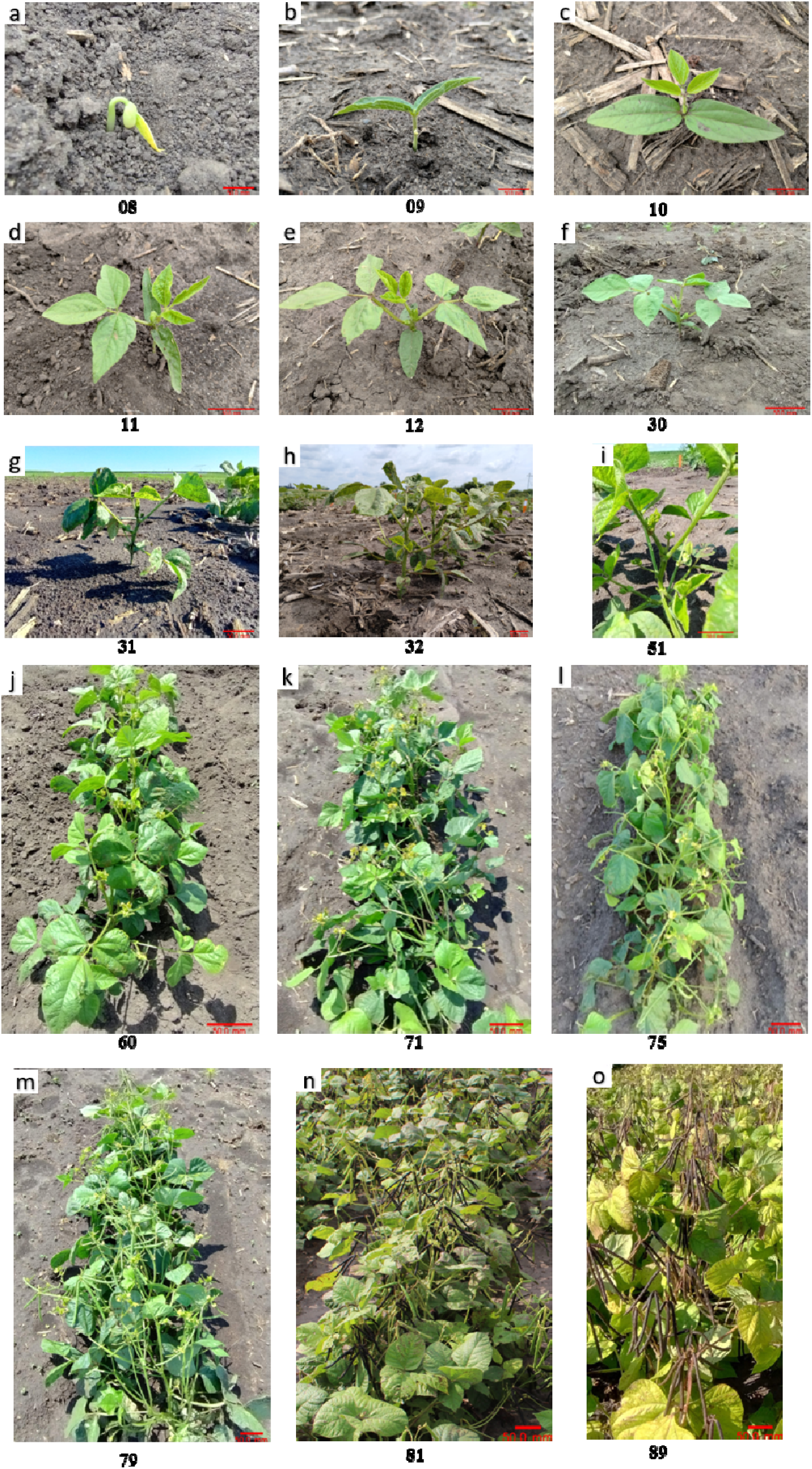
Mungbean key phenological growth stages according to the two-digit BBCH scale: germination (a and b); leaf development (c-e); stem elongation (g and h); flower bud development (i); flowering (j); pod development (k-m); pod maturity (n and o). A pixel to millimeter (mm) scale bar (50 mm) in red color was assigned on the bottom right corner of each image.

**Figure 5.**
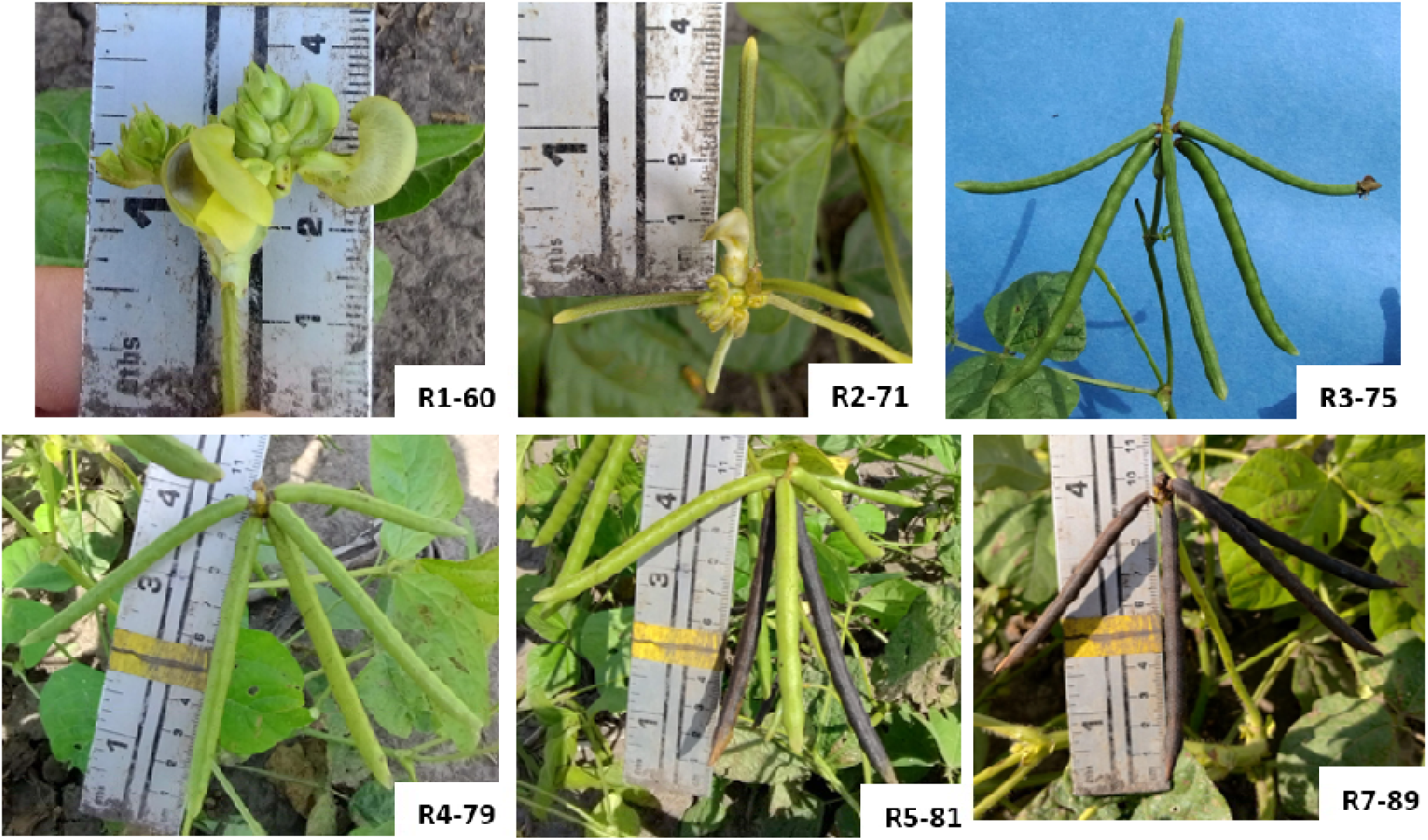
Key mungbean reproductive stages transformation from flowering (R1) to the final maturity (R7) in the ISU Mung G1 cultivar at the Bruner24 environment. Each image i annotated with the V/R growth stage and the BBCH code.

#### BBCH 60: R1 Stage

The first open flower is observed at any of the three upper nodes on the main stem on 50% of the plants in the plot, marking the beginning of flowering, and this stage continues until full bloom. It requires an accumulated thermal unit of 523 °C.d, approximately 42 days after planting.

### 3.6. Principal Growth Stage 7: Pod development

During this stage, the pod initiation begins to continue elongation, and the pod filling continues until the full seed stage (Figure 4 and 5).

#### BBCH 71: R2 Stage

Pod initiation begins with the formation of a pod approximately 1.0 2.0 cm in length and elongates up to 3.0 - 4.0 cm at the end of this stage on the inflorescence between the top three nodes on the main stem. An accumulated thermal unit of 590 °C.d was observed for this stage around 47 days after planting.

#### BBCH 75: R3 Stage

Pod filling begins when a pod approximately 4.0 5.0 cm long is observed at any of the top three nodes on the main stem and continues its elongation till the end of this stage. This stage requires an accumulated thermal unit of 637 °C.d, approximately 52 days after planting.

#### BBCH 79: R4 Stage

At the full seed stage, at least 1-2 pods in a cluster on any of the top three nodes developed constrictions between the seeds. The pods turn pale yellowish or lighter green, indicating they are ready for maturity. This stage occurs with an accumulated thermal unit of 716 °C.d, observed around 57 days after planting.

### 3.7. Principal Growth Stage 8: Ripening (Maturity)

The color of the pods on clusters in any of the top node’s changes from pale yellowish green to brown or black, indicating the beginning of the maturity stage. The percentage of pods that turn black or brown determines each sub-stage.

#### BBCH 81: R5 stage

The first pod in a cluster turns black or brown in color, indicating the beginning of maturity. The total accumulated thermal units for this stage are 800 °C.d. Subsequently, around 50% of the pods on the plant mature 66 days after planting, and 75% at around 69 days after planting.

#### BBCH 88: R6 stage

At this stage, 85-90 % of pods on the plant turn black or brown, indicating near complete maturity to begin harvesting. This stage is reached with an accumulated thermal unit of 908 °C.d, observed at 72 days after planting.

#### BBCH 89: R7 Stage

The plant fully matures when 100% of the pods mature. This stage is reached with an accumulated thermal unit of 983 °C.d, observed approximately at 79 days after planting.

### 3.8 Principal Growth Stage 8: Senescence

We could not observe the senescence stage during these experiments because the two mungbean cultivars we used in this study require chemical desiccation for harvesting. The plants remain green for a longer time after maturity. However, the leaves turn pale yellow with brown spots, and the stems turn brown, eventually dropping when the plants undergo senescence.

**Figure 6:**
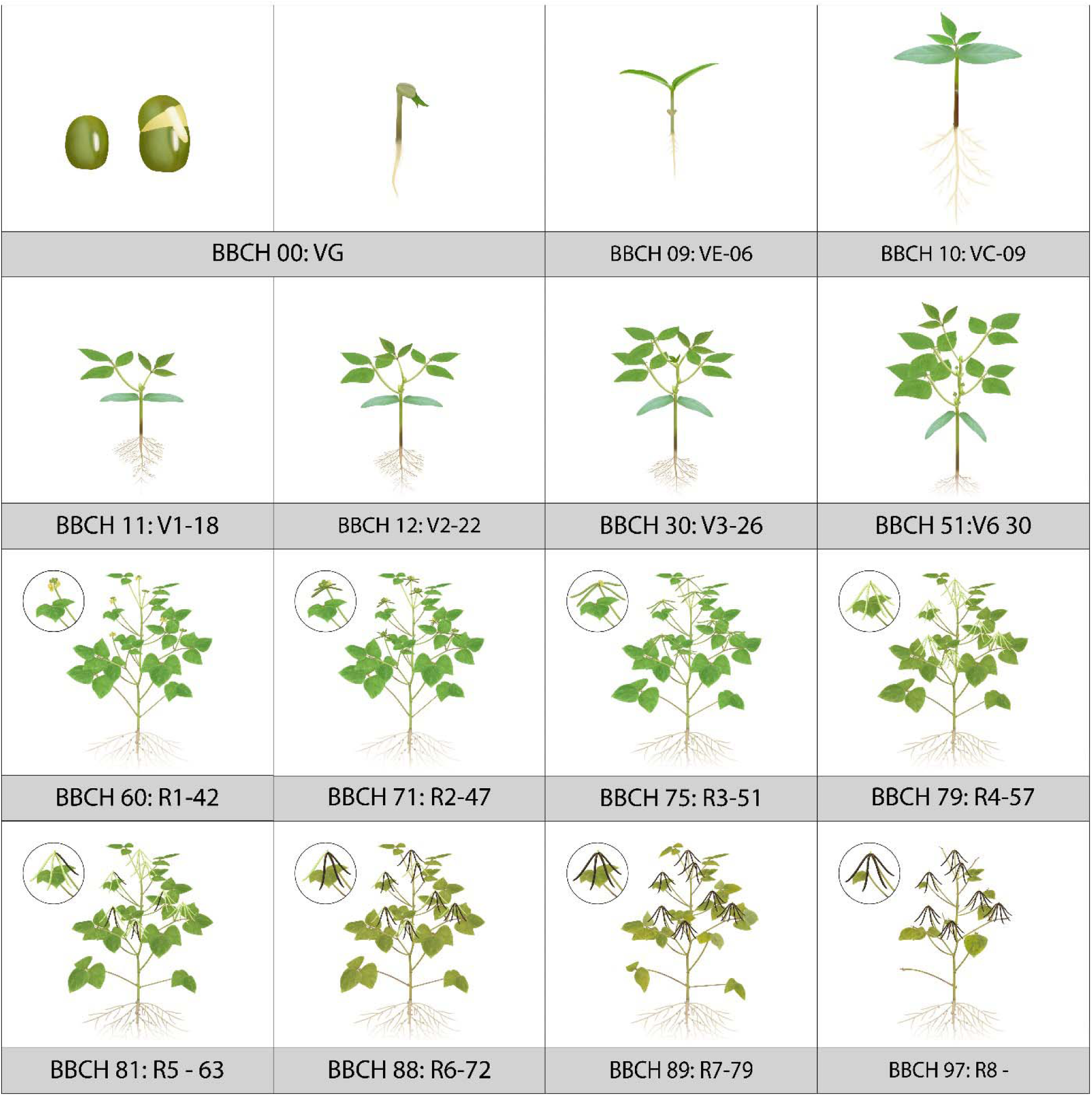
Bio-rendered sketch illustrating Mungbean growth stages according to the two-digit BBCH scale and the V/R stage chart, from germination to senescence.

## 4. DISCUSSION

The two-digit BBCH scale of mungbean phenological stages provides a standardized framework that offers valuable insights into mungbean’s growth and developmental stages, which can be utilized for standardized management practices for yield optimization. This study aimed to delineate the growth stages of mungbean using the BBCH scale and corresponding V/R stages, which is widely recognized for its applicability across various crops (Herraiza et al., 2015; Liu et al., 2015; Acosta-Quezada et al., 2016; Kishore & Mahanti, 2016; Sosa-Zuniga et al., 2017; Ramírez & Kallarackal, 2018; Martínez et al., 2019; Cavalcante et al., 2020; Pati et al., 2020; Pinzón-Sandoval et al., 2024). This is the first BBCH chart developed for mungbean crops globally.

Climate change can affect the duration of the crop cycle and development, pressing the need to understand the relationship between phenological stages and the associated Growing Degree Days (GDD) for each stage. This understanding can help in making necessary adjustments in crop management (Hall et al., 2016; Ramírez & Kallarackal, 2018). One of the primary meteorological factors is air temperature, which influences agricultural crop development (Mendes et al., 2017) and plays a critical role in determining the thermal requirements of crops. The thermal sum is a crucial parameter for reducing climatic risks by measuring the heat units required for each phenological stage to forecast crop cycle duration. Mungbean grows best in temperatures between 25 and 35°C, with optimal growth and yield observed at 28 to 35°C (Kaur et al., 2015; Bhardwaj et al., 2023). Air temperature is a key meteorological factor influencing mungbean productivity in different regions, as extreme temperatures can significantly impact physiological processes and final yield (HanumanthaRao et al., 2016; Malaviarachchi et al., 2016). In addition to air temperature, total rainfall is also one of the key factors affecting mungbean growth. An optimal rainfall distribution for mungbean is 15-20 inches (400-500 mm) (Bhardwaj et al., 2023). During our experiments in 2023 and 2024, we recorded an average of 14 inches (355 mm) of rainfall and an average relative humidity of 78. Integrating the BBCH chart of growth stages with Growing Degree Days (GDD) models allows for the adjustment of planting times, precise irrigation, and crop management decisions to ensure that water and thermal requirements are met at each growth phase. This method can be effective in regions with variable climates, where proactive adjustments can protect productivity.

The BBCH chart can efficiently illustrate the mungbean phenological growth stages, and the two-digit scale can categorize them into principal and secondary stages, providing a clear understanding to growers for monitoring crop growth and development. The principal stages, including germination, vegetative, reproductive, and maturity, were observed to closely align with the mungbean growth patterns in the Midwest region. The secondary stages under each principal stage offer further breakdown, allowing for precise identification of the various growth stages.

The germination stage includes several secondary processes, such as the dry seed reaching the hypocotyl and the soil surface. It begins with seed imbibition, followed by the emergence of the radicle from the seed. Next, the hypocotyl and cotyledons break through the seed coat to reach the soil surface (Cavalcante et al., 2020; Pinzón-Sandoval et al., 2024). The flowering stage (BBCH 60-65) is a critical period in the mungbean growth cycle (Islam et al., 2021). The timing and duration of the flowering impacted the pod set and, ultimately, yield directly. In this study, flowering occurred approximately 40 - 42 days after planting and lasted actively for 5-10 days. This is a critical stage for proper management practices, particularly controlling insects and soil moisture to avoid flower drop due to stress. Also, high temperatures during the flowering stages cause detrimental effects (Gentry, 2010).

Pod development (BBCH 71-79) followed flowering and was characterized by rapid growth and filling of pods. The duration of this stage varies depending on the cultivar and environmental conditions, but generally, pods reach physiological maturity within 20-25 days after flowering. The BBCH scale enabled precise monitoring of pod development, as this stage is particularly vulnerable to pests and diseases. Pesticide spraying schedules are essential to prevent significant crop losses during this stage (Heidt, 2024). The maturation stage (BBCH 81-89) marked the final phase of the mungbean growth cycle. During this stage, the pods turned from pale green to brown or black, indicating the plant transition into maturity. The BBCH scale outlines clearly the maturation process to enable farmers to identify the optimal time for harvest following desiccation or windrowing. Determining the right stage for harvesting is crucial to prevent yield losses due to pod shattering and seed discoloration. In Asian countries, most harvesting is done manually and typically involves at least two rounds of harvesting. The first round occurs at the R6 growth stage, while the second occurs at the R7 stage (Pookpakdi et al., 1992). However, in areas where harvesting is done mechanically, farmers often desiccate the crop at the R6 stage (Dimond, 2024).

This study provides a comprehensive overview of the growth stages of mungbean. However, there are several areas that need further investigation, particularly regarding the effects of climate change on the phenological stages of mungbean in relation to fluctuations in temperature and rainfall patterns. Additionally, examining the relationship between different phenological stages and the occurrence of pests and diseases could aid in developing integrated pest management practices. Applying the BBCH scale to other legumes grown in the Midwest region could improve our understanding of legume phenology. Additionally, conducting comparative studies across various regions may offer valuable insights into the adaptability of mungbean to different agro-ecological conditions.

## 5. CONCLUSION

The mungbean BBCH chart and the two-digit scale, along with the Growing degree days (GDD) developed from this study, could potentially help researchers and growers, particularly in the Midwest region of the United States. This chart can be useful in designing better mungbean management practices for enhancing its production. The demand for plant-based proteins continues to rise, and the role of legumes like mungbean significantly increases. It presses the need to understand the growth and development of mungbean will be increasingly essential to cater to the needs of this plant-based protein industry.

## ACKNOWLEDGMENT

This work was supported by funding from the United States Department of Agriculture - National Institute of Food and Agriculture (USDA-NIFA) under Mung bean breeding #2022–67013-37120, the RF Baker Center for Plant Breeding, and the department of Agronomy. Additional support was provided by the United States Department of Agriculture Hatch CRIS project IOW04714.

## CONFLICT OF INTEREST

The authors state they have no conflicts of interest related to this manuscript’s submission and publication.

## Notes

### Competing Interest Statement

The authors have declared no competing interest.

### Summary of Updates

This version fixed the labelling errors in the figure 6, BBCH skecth chart.

