## Supplemental Table 1-3 (Table S1-S3) for "Defining the Growth Stages of Mungbean (*Vigna radiata* L.) using the BBCH Scale" (from another version of this preprint: version 1, 2025-11-11)

**Table S1:** Weekly weather data on mean maximum temperature, mean minimum temperature, precipitation, relative humidity, and growing degree days (GDD) of each site-year during the experimental duration.

| **site_year** | **week** | **max_temp (c)_mean** | **min_temp (c)_mean** | **precip (inch) _sum** | **avg(RH) _mean** | **GDD_ C_sum** |
| --- | --- | --- | --- | --- | --- | --- |
| aea23 | 1 | 29.52 | 17.97 | 0.35 | 73.6 | 96.22 |
| aea23 | 2 | 26.57 | 12.55 | 0 | 56.41 | 66.92 |
| aea23 | 3 | 29.11 | 16.06 | 1.66 | 68.96 | 88.1 |
| aea23 | 4 | 28.14 | 17.37 | 1.4 | 75.36 | 89.28 |
| aea23 | 5 | 28.71 | 18.55 | 0.56 | 82.78 | 95.38 |
| aea23 | 6 | 25.5 | 13.9 | 3.61 | 79.2 | 67.91 |
| aea23 | 7 | 26.89 | 15.79 | 0 | 83.66 | 79.4 |
| aea23 | 8 | 29.97 | 17.27 | 0.25 | 80.86 | 95.34 |
| aea23 | 9 | 29.69 | 19.08 | 0.48 | 86.08 | 100.72 |
| aea23 | 10 | 27.66 | 18.35 | 3.41 | 87.72 | 91.03 |
| aea23 | 11 | 27.43 | 15.44 | 0.57 | 84.54 | 80.05 |
| aea23 | 12 | 30.93 | 18.97 | 0 | 85.06 | 104.66 |
| aea23 | 13 | 29.06 | 17.17 | 0.03 | 79.51 | 91.8 |
| aea23 | 14 | 31.01 | 14.61 | 0 | 73.94 | 89.68 |
| aea23 | 15 | 25.61 | 11.68 | 0.27 | 73.55 | 60.51 |
| aea23 | 16 | 25.12 | 11.19 | 1.32 | 76.37 | 57.07 |
| aea23 | 17 | 24.57 | 13.79 | 0.01 | 85.91 | 64.26 |
| aea23 | 18 | 29.97 | 14.91 | 0 | 75.27 | 37.32 |
| aea24 | 1 | 28.33 | 15.51 | 1.06 | 69.08 | 83.44 |
| aea24 | 2 | 28.87 | 15.53 | 0 | 55.18 | 85.42 |
| aea24 | 3 | 29.36 | 18.63 | 1.47 | 79.97 | 97.95 |
| aea24 | 4 | 30.01 | 19.43 | 0.67 | 78.76 | 103.01 |
| aea24 | 5 | 25.46 | 15.16 | 0.83 | 81.5 | 72.18 |
| aea24 | 6 | 27.93 | 16.47 | 0.97 | 81.07 | 85.38 |
| aea24 | 7 | 29 | 18.23 | 0.19 | 81.33 | 95.31 |
| aea24 | 8 | 27.57 | 18.1 | 0.31 | 88.73 | 89.84 |
| aea24 | 9 | 30.06 | 19.5 | 1.17 | 85.63 | 103.45 |
| aea24 | 10 | 27.91 | 16.75 | 0 | 81.2 | 86.32 |
| aea24 | 11 | 24.79 | 15.7 | 0.72 | 87.45 | 71.74 |
| aea24 | 12 | 25.77 | 14.87 | 0.11 | 82.48 | 72.22 |
| aea24 | 13 | 31.03 | 19.42 | 0.89 | 85.56 | 106.57 |
| aea24 | 14 | 25.16 | 12.17 | 0 | 74.56 | 60.66 |
| aea24 | 15 | 26.72 | 11.73 | 0 | 69.25 | 64.57 |
| aea24 | 16 | 30.73 | 17.35 | 1.19 | 76.43 | 98.28 |
| aea24 | 17 | 25.96 | 11.29 | 0.58 | 71.02 | 60.37 |
| aea24 | 18 | 29.75 | 13.48 | 0 | 63.14 | 34.84 |
| bruner24 | 1 | 27.7 | 15.33 | 1.18 | 69.26 | 80.62 |
| bruner24 | 2 | 28.19 | 14.32 | 0 | 58.45 | 78.77 |
| bruner24 | 3 | 29.03 | 19.16 | 1.8 | 77.88 | 98.68 |
| bruner24 | 4 | 29.26 | 19.02 | 1.31 | 79.39 | 98.97 |
| bruner24 | 5 | 24.94 | 14.66 | 1.07 | 82.08 | 68.61 |
| bruner24 | 6 | 27.43 | 15.31 | 1.55 | 83.35 | 79.59 |
| bruner24 | 7 | 28.16 | 17.04 | 0.41 | 83.88 | 88.2 |
| bruner24 | 8 | 27.42 | 17.47 | 0.91 | 88.47 | 87.14 |
| bruner24 | 9 | 29.4 | 19.27 | 1.25 | 85.8 | 100.38 |
| bruner24 | 10 | 27.51 | 15.71 | 0 | 82.18 | 81.3 |
| bruner24 | 11 | 24.53 | 14.9 | 1 | 87.91 | 68 |
| bruner24 | 12 | 25.43 | 13.63 | 0.21 | 83.56 | 66.7 |
| bruner24 | 13 | 30.75 | 18.57 | 1.36 | 85.44 | 102.62 |
| bruner24 | 14 | 25.11 | 10.79 | 0 | 77.31 | 55.67 |
| bruner24 | 15 | 26.46 | 10.05 | 0 | 71.55 | 57.78 |
| bruner24 | 16 | 30.2 | 16.28 | 1.37 | 76.63 | 92.66 |
| bruner24 | 17 | 25.45 | 9.49 | 0.76 | 76.36 | 52.31 |
| bruner24 | 18 | 29.91 | 10.91 | 0 | 67.79 | 31.24 |

**Table S2:** Raw data on each growth stage of two cultivars from three environments tested in 2023 and 2024**.**

| **env** | **block** | **cultivar** | **PS0-VE** | **PS1-VC** | **PS1-V1** | **PS1-V2** | **PS3-V3** | **PS3-V4** | **PS3-V5** | **PS5-V6** | **PS6-R1** | **PS7-R2** | **PS7-R3** | **PS7-R4** | **PS8-R5** | **PS8-R6** | **PS8-R7** |
| --- | --- | --- | --- | --- | --- | --- | --- | --- | --- | --- | --- | --- | --- | --- | --- | --- | --- |
| aea23 | 1 | ISU Mung G-1 | 6 | 13 | 20 | 26 | 32 | 35 |  |  | 47 | 55 | 62 | 69 | 76 | 81 | 85 |
| aea23 | 1 | ISU Mung G-2 | 7 | 14 | 21 | 28 | 32 | 35 | 42 | 47 | 49 | 51 | 59 | 62 | 69 | 76 | 81 |
| aea23 | 2 | ISU Mung G-1 | 7 | 14 | 21 | 25 | 28 | 32 | 35 |  | 47 | 51 |  |  |  |  |  |
| aea23 | 2 | ISU Mung G-2 | 7 | 14 | 21 | 28 | 32 | 35 | 42 | 45 | 47 | 51 | 55 | 59 | 62 | 69 | 76 |
| aea23 | 3 | ISU Mung G-1 | 7 | 14 | 21 | 25 | 28 | 32 | 35 | 42 | 47 | 51 |  |  |  |  |  |
| aea23 | 3 | ISU Mung G-2 | 7 | 14 | 21 | 28 | 35 | 47 | 51 |  |  |  |  |  |  |  |  |
| aea23 | 4 | ISU Mung G-1 |  | 14 | 21 | 28 | 32 | 35 | 42 |  | 47 | 51 | 55 | 59 |  |  |  |
| aea23 | 4 | ISU Mung G-2 | 7 | 14 | 21 | 28 | 32 | 32 | 35 | 42 | 51 | 55 |  |  |  |  |  |
| aea23 | 5 | ISU Mung G-1 | 5 | 12 | 18 | 25 | 32 | 35 | 42 |  | 47 | 51 | 55 | 59 | 62 | 69 | 76 |
| aea23 | 5 | ISU Mung G-2 |  | 14 | 21 | 28 | 32 | 35 | 42 | 47 | 47 | 51 | 55 | 59 | 69 | 76 | 81 |
| aea24 | 1 | ISU Mung G-1 | 6 | 9 | 15 | 18 | 22 | 29 | 33 | 36 | 36 | 40 | 46 | 51 | 57 | 67 | 76 |
| aea24 | 1 | ISU Mung G-2 | 5 | 8 | 15 | 19 | 23 | 28 | 31 | 35 | 40 | 47 | 52 | 58 | 63 | 73 | 78 |
| aea24 | 2 | ISU Mung G-1 | 5 | 8 | 15 | 18 | 22 | 25 | 30 | 33 | 36 | 40 | 45 | 52 | 58 | 65 | 73 |
| aea24 | 2 | ISU Mung G-2 | 4 | 8 | 15 | 20 | 23 | 26 | 30 | 35 | 40 | 44 | 52 | 58 | 63 | 70 | 78 |
| aea24 | 3 | ISU Mung G-1 | 5 | 8 | 15 | 18 | 22 | 27 | 31 | 34 | 38 | 42 | 46 | 53 | 58 | 68 | 76 |
| aea24 | 3 | ISU Mung G-2 | 6 | 8 | 15 | 19 | 23 | 27 | 31 | 34 | 40 | 44 | 52 | 57 | 63 | 73 | 78 |
| aea24 | 4 | ISU Mung G-1 | 5 | 8 | 15 | 18 | 22 | 26 | 31 | 34 | 38 | 44 | 49 | 54 | 60 | 73 | 79 |
| aea24 | 4 | ISU Mung G-2 | 4 | 8 | 15 | 19 | 23 | 26 | 30 | 33 | 40 | 44 | 50 | 56 | 59 | 68 | 78 |
| aea24 | 5 | ISU Mung G-1 | 5 | 8 | 17 | 19 | 23 | 26 | 30 | 34 | 37 | 44 | 49 | 54 | 59 | 69 | 76 |
| aea24 | 5 | ISU Mung G-2 | 5 | 8 | 17 | 20 | 23 | 26 | 30 | 34 | 40 | 45 | 50 | 57 | 63 | 73 | 78 |
| bruner24 | 1 | ISU Mung G-1 | 6 | 9 | 16 | 20 | 24 | 27 | 33 | 36 | 39 | 45 | 50 | 56 | 63 | 73 | 78 |
| bruner24 | 1 | ISU Mung G-2 | 6 | 9 | 17 | 20 | 25 | 28 | 32 | 36 | 41 | 46 | 52 | 58 | 63 | 74 | 81 |
| bruner24 | 2 | ISU Mung G-1 | 7 | 11 | 16 | 19 | 21 | 25 | 31 | 36 | 38 | 44 | 48 | 56 | 60 | 71 | 78 |
| bruner24 | 2 | ISU Mung G-2 | 7 | 10 | 16 | 19 | 22 | 25 | 32 | 37 | 40 | 45 | 49 | 58 | 63 | 71 | 80 |
| bruner24 | 3 | ISU Mung G-1 | 8 | 11 | 15 | 19 | 21 | 25 | 30 | 35 | 35 | 38 | 43 | 50 | 57 | 71 | 78 |
| bruner24 | 3 | ISU Mung G-2 | 6 | 10 | 16 | 21 | 23 | 27 | 31 | 36 | 39 | 44 | 48 | 56 | 62 | 73 | 80 |
| bruner24 | 4 | ISU Mung G-1 | 6 | 11 | 16 | 19 | 21 | 25 | 31 | 36 | 39 | 45 | 49 | 54 | 61 | 72 | 78 |
| bruner24 | 4 | ISU Mung G-2 | 5 | 9 | 15 | 19 | 21 | 25 | 29 | 33 | 40 | 47 | 50 | 56 | 63 | 74 | 80 |
| bruner24 | 5 | ISU Mung G-1 | 6 | 11 | 18 | 20 | 23 | 27 | 30 | 35 | 39 | 45 | 48 | 52 | 59 | 71 | 79 |
| bruner24 | 5 | ISU Mung G-2 | 5 | 10 | 17 | 19 | 21 | 25 | 29 | 35 | 40 | 46 | 52 | 58 | 73 | 78 | 85 |

**Table S3**: The results of the study on the phenological growth stages of the mungbean based on the two-digit BBCH scale and corresponding V/R stage codes.

| **BBCH_Code** | **V/R_Stage** | **Stage Description** |
| --- | --- | --- |
| **Principal growth stage 0: Germination** | | |
| 00 |  | Dry seed |
| 01 |  | Seed imbibition beginning |
| 03 |  | Seed imbibition complete |
| 05 |  | Radicle emergence from the seed |
| 07 |  | Hypocotyl with cotyledons emerged from the seed coat |
| 08 | VG - Germination | Hypocotyl reaches the soil surface with a visible arch |
| 09 | VE - Emergence | Emergence: Cotyledons with plumule emerged from the soil surface |
| **Principal growth stage 1: Leaf development** | | |
| 10 | VC - Cotyledon | Cotyledons are fully expanded and unifoliate leaves above unroll so that the leaves are not touching each other. |
| 11 | V1 - First trifoliate leaf | Unifoliate leaves attached to the first are fully expanded and flat, while the first trifoliate leaf attached to the upper node starts unrolling |
| 12 | V2 - Second trifoliate leaf | The first trifoliate leaf attached opposite to the second node is fully expanded and flat, while the second trifoliate leaf attached to the upper node starts unrolling. |
| **Principal growth stage 3: Stem elongation** | | |
| 30 | V3 - Third trifoliate leaf | Stem elongation begins with the second trifoliate leaf attached to the third node is fully expanded and flat, while the third trifoliate leaf attached to the upper node starts unrolling. |
| 31 | V4 - Fourth trifoliate | First visible internode extension coupled with the third trifoliate leaf attached to the fourth node is fully expanded and flat, while the fourth trifoliate leaf attached to the second upper node starts unrolling. |
| 32 | V5 - Fifth trifoliate | The second visible internode extension, along with the fourth trifoliate leaf attached to the fourth node, is fully expanded and flat, while the fourth trifoliate leaf attached to the second upper node starts unrolling. |
| 33 | V6 - Sixth trifoliate | The fifth trifoliate leaf attached to the sixth node is fully expanded and flat, while the sixth trifoliate leaf attached to the upper node starts unrolling. |
| **Principal growth stage 6: Flowering (Reproductive stage)** | | |
| 60 | R1 - Flowering | First open flower at the uppermost flowering nodes |
| 65 |  | Open flowers at two or more uppermost nodes |
| **Principal growth stage 7: Development of fruit (Pod maturity)** | | |
| 71 | R2 - Pod initiation | First pod 2-5 cm long at one of four uppermost nodes |
| 75 | R3 - Pod filling | 50% of the pods on the plants reached maturity, and the seeds began to fill out. |
| 79 | R4 - Full Seed | up to 90% of the pods having easily visible individual beans with the constrictions on the pod surface |
| **Principal growth stage 8: Ripening of fruit and seed (Pod and Seed maturity)** | | |
| 81 | R5 - Beginning seed maturity | Pods contain full-size seeds - beginning maturity with around 10% of matured pods |
| 82 |  | 25% of the pods on the plants turn black/brown |
| 85 |  | 50% of the pods on the plants turn black/brown |
| 87 |  | 75% of the pods on the plants turn black/brown |
| 88 | R6 - Harvest maturity | 90% of the Pods contain full-sized hard seeds |
| 89 | R7 - Final maturity | Full maturity: Pods changed color with fully matured seed |
| **Principal growth stage 9: Senescence** | | |
| 97 | R8 - Senescence | Senescence: leaves dried and dropped; plant dead. |
| 99 |  | Harvested grains |
